# Genetic mechanisms for tissue-specific essential genes

**DOI:** 10.1101/2021.04.09.438977

**Authors:** Elad Dvir, Shahar Shohat, Sagiv Shifman

**Affiliations:** Department of Genetics, The Institute of Life Sciences, The Hebrew University of Jerusalem, Jerusalem, Israel

## Abstract

Genetic diseases often manifest in specific tissues despite having the genetic risk variants in all cells. The most commonly assumed mechanism is selective expression of the causal gene in the pathogenic tissues, but other mechanisms are less explored. Using CRISPR screens from 789 cell lines and 27 lineages, we identified 1274 lineage-specific essential genes (LSEGs). We show that only a minority of LSEGs are explained by preferential expression (n = 115), and a big proportion of them (n = 509) is explained by lineage-specific gene amplification. Three other mechanisms were identified by genome-wide expression analysis. First, lineage-specific expression of paralogs leads to reduced functional redundancy and can account for 153 LSEGs. Second, for 45 LSEGs, the paralog expression increases vulnerability, implying functional codependency. Third, we suggest that the transfer of small molecules to mutant cells could explain blood-specific essentiality. Overall, LSEGs were more likely to be associated with human diseases than common essential genes, were highly intolerant to mutations and function in developmental pathways. Analysis of diverse human cell types found that the expression specificity of LSEGs and their paralogs is consistent with preferential expression and functional redundancy being a general phenomenon. Our findings offer important insights into genetic mechanisms for tissue specificity of human diseases.

## Introduction

The consequence of a mutation depends on the genetic and environmental context of the cells or organism ^1^. Thus, a loss-of-function mutation may result in a phenotype in a particular cell type that is more vulnerable to losing this gene ^2^. For example, hereditary diseases, including neurodegenerative disorders, cardiovascular diseases, and autoimmune diseases, often affect a specific tissue, although the genetic risk variants are present across the human body ^2–4^. Similarly, some genes are known to be essential during specific developmental processes. For example, it was found that during mammalian cardiac development, there is a functional diversity of essential genes, such as some genes are crucial in the early stages and others in more progress stages ^5^.

Preferential expression or even exclusive expression in the vulnerable cells may suggest specific gene functions in those cells ^2,6,7^. For example, the caveolin 3 (CAV3) gene is selectively expressed in cardiac myocytes, skeletal muscle, and smooth muscle cells, and its mutation causes cardiomyopathies and skeletal muscle disorders ^8^. Indeed, a large-scale analysis found that disease genes and complexes tend to be overexpressed in tissues where mutations cause pathology ^6^. It has also been shown that genes up-regulated in cancer cell lines are more essential in cancer cells than non-cancer cells ^9,10^.

The development of CRISPR-Cas9 based approaches for genome-wide loss-of-function (LoF) screening made it possible to identify essential genes in large panels of cell lines ^11,12^. Project Achilles used a large panel of human cancer cell lines representing multiple cancer lineages to create a catalog of essential genes ^13^. The fitness consequences of disrupting a specific gene are estimated by comparing the abundances of single-guide RNAs (sgRNAs) targeting the gene at the start of the experiment to the levels of the same sgRNAs after a few weeks of growth represented by a dependency score. A relative decrease in the abundance of sgRNAs suggests that disruption of the target gene causes a viability defect. This resource makes it possible to systematically investigate which genes are essential for cancer cells and discover the factors that contribute to the variation in essentiality.

In the current study, we utilized the large-scale CRISPR screens to identify lineage-specific essential genes (LSEGs) and study mechanisms underlying their specificity. We identified 1274 unique LSEGs, and we characterized which genes are best explained by preferential expression, functional redundancy, or functional codependency based on gene expression from the same cells. Additionally, we found two other mechanisms for LSEGs. For some genes, gene amplification was associated with increased gene expression and decreased essentiality, probably due to insufficient disruption of all gene copies. For genes essential specifically in blood cells, we suggest that intracellular communication with an exchange of small metabolites between cells, a mechanism that is absent in blood cells, can explain the lineage specificity. We show that LSEGs are intolerant to functional mutations and are especially associated with human diseases, suggesting the general importance of LSEGs. We also observed that LSEGs and their paralogs show different cell-type-specific expression patterns depending on the suggested mechanism, consistent with those mechanisms being conserved across cell types.

## Results

### Inferring lineage-specific essential genes from large-scale CRISPR screens

To comprehensively catalog genes essential for specific lineages, we utilized data from large-scale CRISPR screens in cancer cell lines to identify genes required for growth or viability (789 cancer cell lines originated from 27 different lineages). The dataset contains a dependency score for each gene in each cell line, and a very negative dependency score implies that the cell line is highly dependent on that gene. The scores are adjusted such as a score of 0 is equivalent to a gene under no negative selection, and the median score of the highly essential core genes is -1. Lineage-specific essential genes (LSEGs) were defined as genes with significantly lower levels of dependency scores in a specific lineage of cells compared to all other lineages (FDR < 0.05) and with an average dependency score < -0.5. Using this method, we were able to identify genes that show lineage-specificity even if it is in more than one lineage.

We identified 1701 LSEGs in 23 lineages, including 1274 unique genes (Table S1). The number of genes per lineage varied, with the highest numbers for plasma and blood (Figure 1A). The number of cell lines per lineage is unlikely to explain this variation, as there was no significant correlation between sample size and the number of genes identified per lineage (r = -0.048, *P* = 0.83) (Figure 1B). It is important to note that the dependency scores of LSEG were continuous across lineages for some genes (Figure 1C). In contrast, there was a clear distinction between lineages with low and high dependency scores for other genes (Figure 1D). Since some of the LSEGs were shared between lineages, we quantified the percentage of overlap between lineages (Figure 1E). The highest overlap was found between related lineages, such as blood/plasma, plasma/lymphocytes, blood/lymphocytes, ovary/uterus and pancreas/bile duct (Jaccard Index = 0.186, 0.095, 0.093, 0.091, 0.066, respectively).

**Figure 1.**
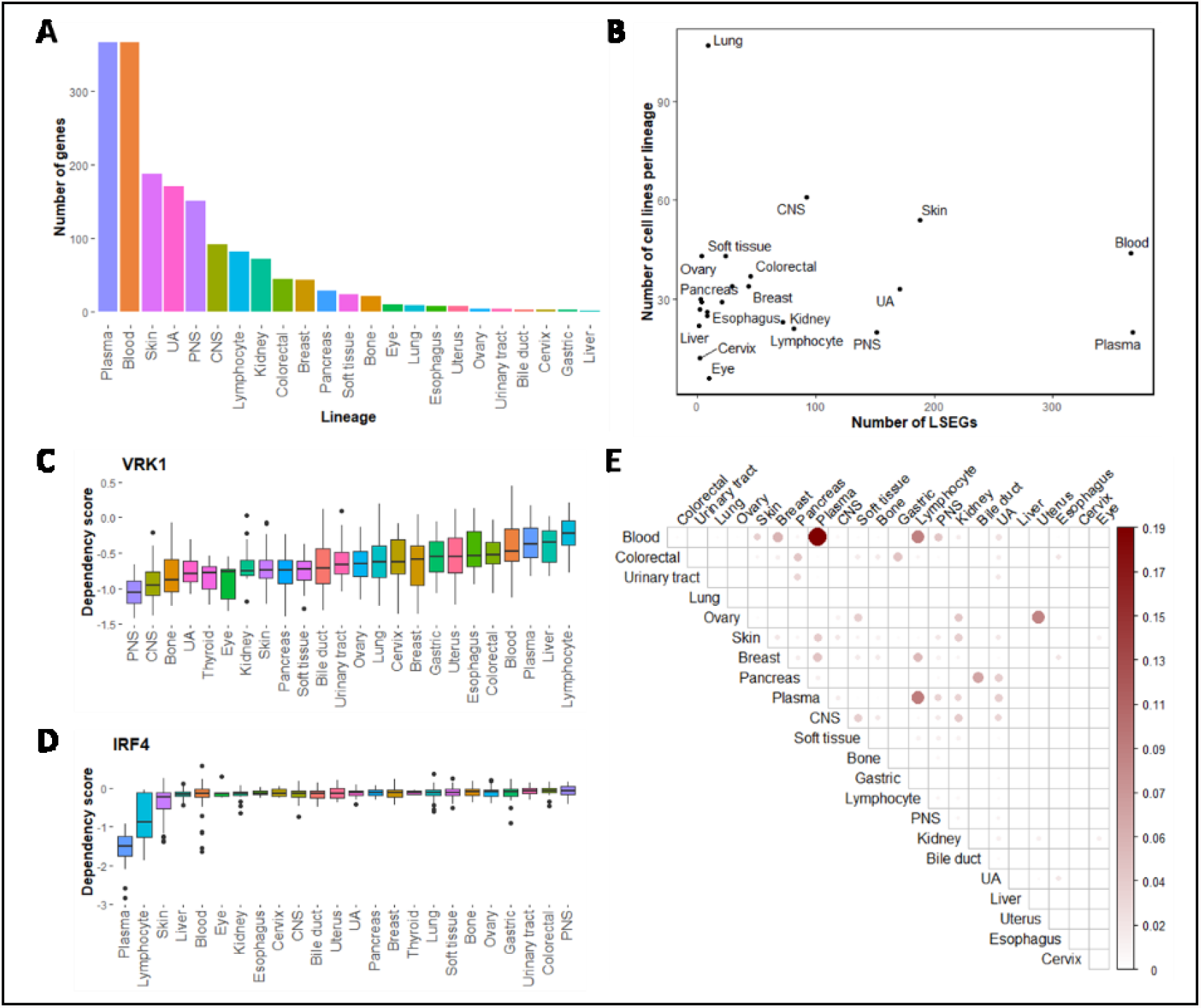
Identification and characterization of lineage-specific essential genes (LSEGs). (A) The number LSEGs per lineage. (B) Number of LSEGs per lineage as a function of sample size (number per lineage). (C-D) Examples of distribution of dependency scores across lineages for two LSEGs. (C) VRK1 is essential in the central and peripheral nervous system, and (D) IRF4 is essential in the plasma and lymphocytes. (E) The overlap of LSEGs between lineages. The overlap was quantified by a metric of intersection over union, also known as the Jaccard index. Abbreviations: CNS – *central nervous system*, PNS – *peripheral nervous system, UA – upper aerodigestive*.

### Analysis of preferential expression and copy number variations suggests that lineage-specific functional processes can explain LSEGs

The simplest explanation of LSEGs is that some genes have restricted function in specific lineages. Those genes with restricted function are expected to show preferential or exclusive expression in the susceptible lineages. To explore this possibility, we used the expression data available for the same cell lines to test for differential expression of 19,144 genes between the lineages with low dependency scores (hereafter the vulnerable lineages) and all other lineages (hereafter the non-vulnerable lineages). While in many cases, the average expression of LSEGs in the vulnerable lineages was considerably higher than the non-vulnerable lineages, surprisingly, the expression of many other LSEGs was lower in the vulnerable lineages (Figure 2A). The tendency for more extreme expression values (lower or higher) in the vulnerable lineages was significantly different than expected by chance based on a randomization test (*P* = 9.9×10^-5^) (Figure 2B).

**Figure 2.**
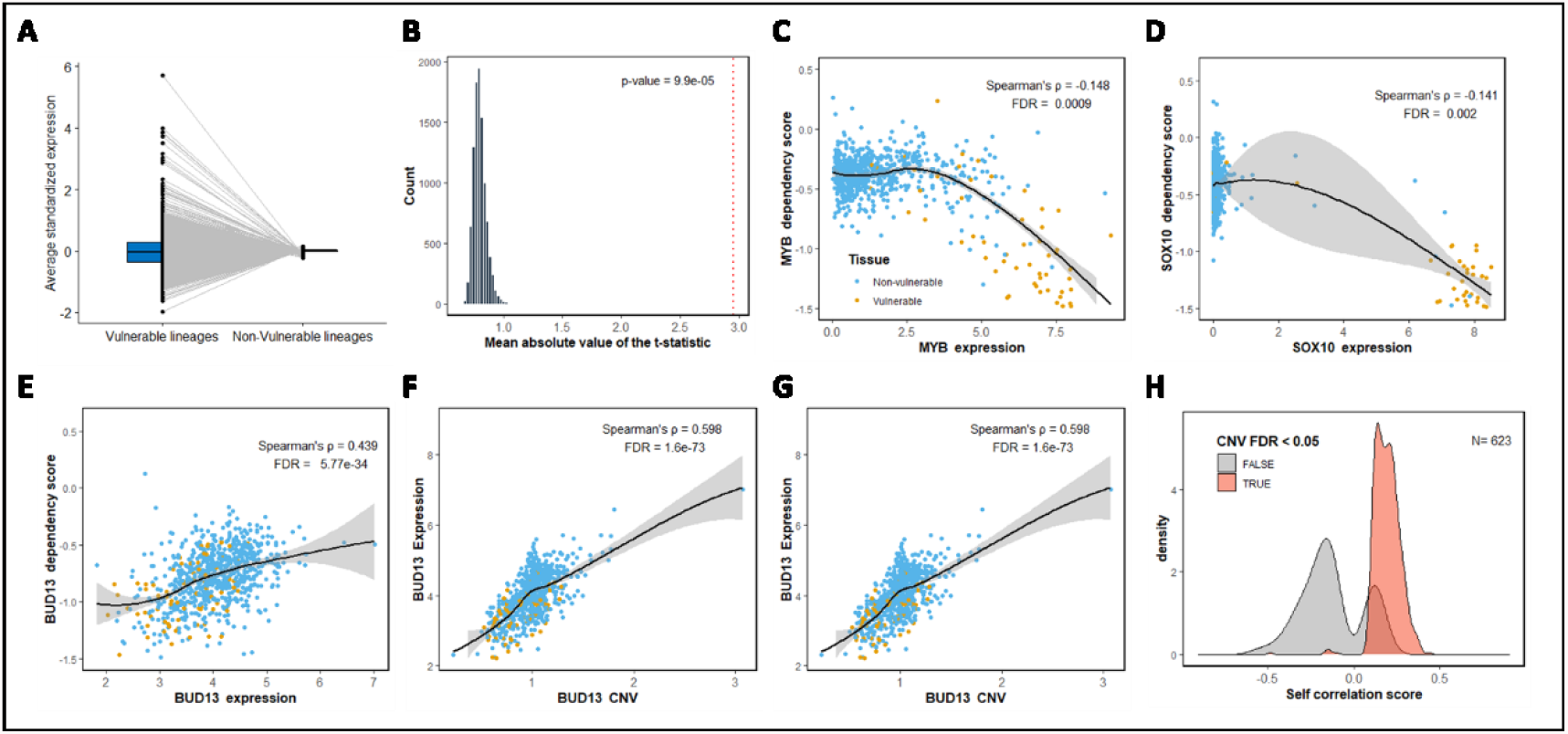
The increase and decrease expression of the LSEGs in vulnerable lineages is consistent with a preferential expression mechanism and the effect of copy number variations (CNVs). (A) The average standardized expression of LSEGs in vulnerable and non-vulnerable lineages. (B) The observed mean absolute t-statistic (dashed red line) was calculated for differences in LSEGs expression between the vulnerable and non-vulnerable lineages. The histogram shows the distribution of the expected values under the null based on 10,000 permutations. (C-D) Demonstration of preferential expression. (C) *MYB*, an LSEG in blood and lymphocytes, is more essential (lower dependency) in cells with higher expression of *MYB*. (D) *SOX10*, an LSEG in the skin, is more essential in cells with higher expression of *SOX10*. (E-G) Demonstration of LSEG that is explained by the gene copy number. (E) *BUD13*, an LSEG in the skin and the upper aerodigestive tissue, is less essential in cells with higher expression of *BUD13*. (F) *BUD13* expression is associated with the copy number of *BUD13*. (G) *BUD13* dependency scores are associated with the copy number of *BUD13*. (H) Density plots of Spearman’s correlation coefficients between the dependency scores and LSEGs expression. The plots are shown for 623 LSEGs with CNV information, divided into genes with significant correlation with the gene copy number (FDR < 0.05) (light red) or non-significant (gray). All *P*-values presented in the figures are adjusted for multiple tests by the Benjamini and Hochberg false discovery rate (FDR) procedure. The yellow dots (C-G plots) represent cell lines that belong to the vulnerable lineages, and blue dots are for cell lines in the non-vulnerable lineages. In C-G, locally estimated scatterplot smoothing (LOESS) curves and 95% confidence intervals are shown alongside Spearman’s ρ and the FDR value.

To estimate how many LSEGs are likely to be explained by preferential expression, we tested for each LSEG the correlation between the dependency scores and the expression levels for all available genes. For the genes that are explained by a preferential expression, we expected that the dependency scores of the LSEG would be negatively correlated with the self-expression of the LSEG (higher expression of the LSEG in cell lines with a lower score that indicates its higher essentiality). Out of the 1274 LSEGs, 624 (49%) had a significant correlation between the dependency scores and their expression (FDR < 0.05), however, only 115 of them were in the direction consistent with preferential expression in the vulnerable lineage (Spearman’s ρ < 0, FDR < 0.05) (see examples in Figure 2C-D).

The remaining 509 LSEGs with a positive correlation (Spearman’s ρ > 0, FDR < 0.05) are genes that are less essential in cell lines with higher expression (see example in Figure 2E). One possible explanation for this unexpected finding is an increase in the copy number of those genes in specific lineages. The multiple copies of the gene can increase the gene expression and decrease the likelihood of a complete knockout by CRISPR. Indeed, we found that that for 469 genes out of the 509 LSEGs (92.1%), there was a significant (FDR < 0.05) positive correlation between the copy number variation (CNVs) and the dependency score (see examples in Figure 2E-G). This association was not common for the 115 LSEGs with preferential expression in the vulnerable lineage, as only 5 (4.3%) showed a significant correlation between the dependency scores and the copy numbers. Overall, the correlation direction between the dependency scores and the gene expression (negative vs. positive) was highly significantly associated with CNVs as predictors (odds ratio (OR) = 250.7, *P* = 2.7×10^-80^) (Figure 2H).

These analyses suggest that a restricted function in specific lineages indicated by preferential expression may explain only a minority of LSEGs (9%). However, gene amplification associated with overexpression of the amplified gene, which occurs more frequently in specific lineages, may explain a considerable proportion (36.8%) of the LSEGs.

### Lineage-specific expression of paralogs may explain a large proportion of LSEGs

Another possible mechanism for the variable gene essentiality between lineages is the expression variation of other genes that can modify the effect of the mutation. We tested the possibility that lineage-specific functional redundancy, which usually involves paralogs, can compensate for the mutated gene. Based on this suggestion, we predicted a lower expression in the vulnerable lineages of the paralogs. To study the expression of paralogs as a potential mechanism, we first identified LSEGs with paralogs (Table S2). For most LSEGs (n = 867), we did not find any paralogs, and out of the 407 LSEGs with paralogs, most had a small number of paralogs (mode = 1, median = 2) (Figure 3A). Next, we compared the expression of the paralogs in the vulnerable vs. non-vulnerable lineages for 404 LSEGs that had expression data for at least one identified paralog. Similar to the results of the self-expression analysis above, we observed that the average expression of the paralogs tends to be either higher or lower in the vulnerable lineages relative to the non-vulnerable lineages (Figure 3B). The absolute differences in the expression of paralogs between the vulnerable and non-vulnerable lineages were significantly different than expected by chance based on a randomization test (*P* = 9.9×10^-5^) (Figure 3C).

**Figure 3.**
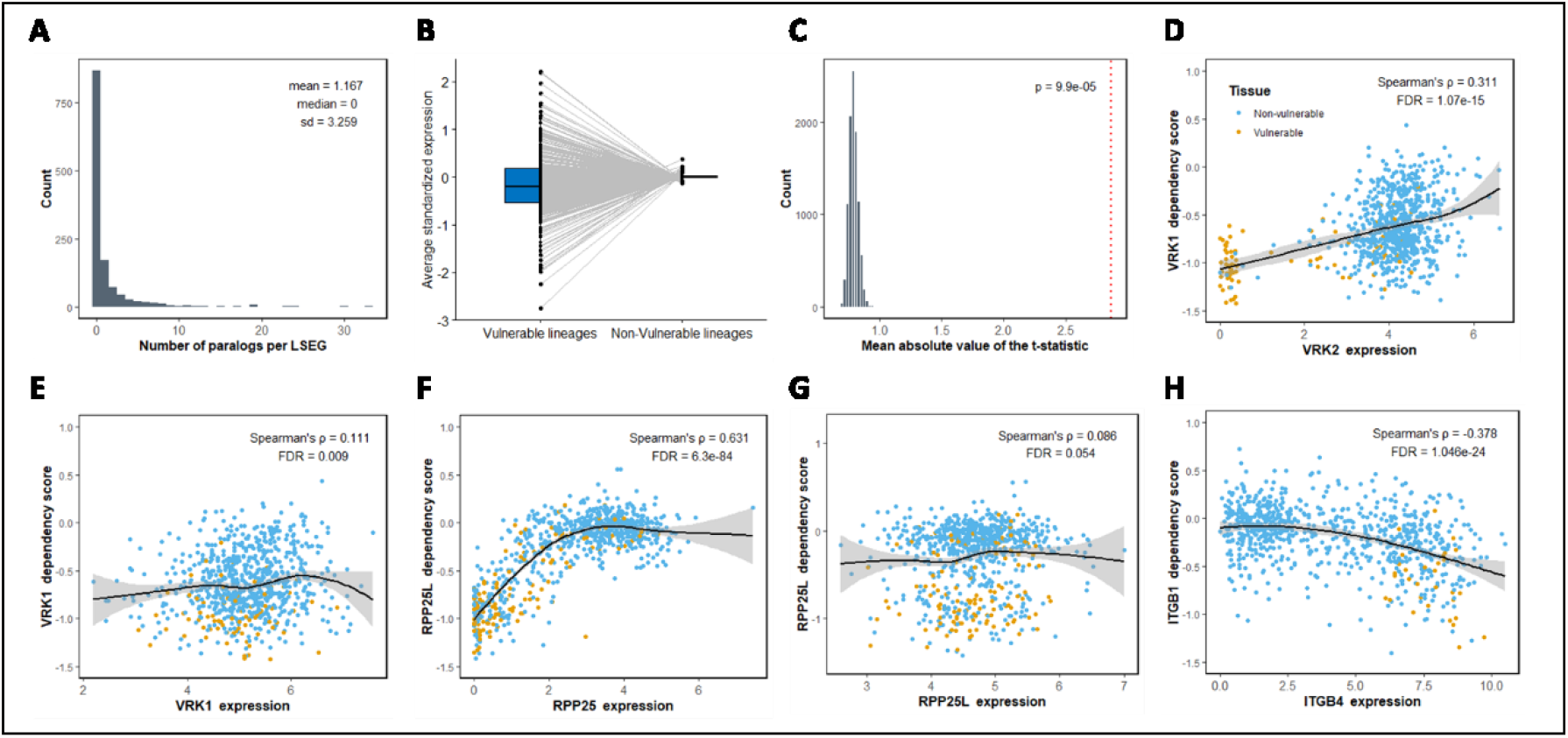
Analysis of paralogs expression suggests functional redundancy and codependency mechanisms for LSEGs. (A) Histogram showing the number of paralogs per each LSEG. (B) The average standardized expression of LSEGs paralogs in vulnerable and non-vulnerable lineages. (C) The observed mean absolute t-statistic (dashed red line) was calculated for differences in paralog expression between the vulnerable and non-vulnerable lineages. The histogram shows the distribution of the expected values under the null based on 10,000 permutations. (D-G) Demonstration of functional redundancy. (D-E) The dependency of *VRK1*, an LSEG in the central and peripheral nervous systems, is correlated mainly with (C) the expression of its paralog, *VRK2*, (D) relative to the expression of *VRK1* itself. (F-G) The dependency of RPP25L, an LSEG in the central nervous systems, soft tissues, and lymphocytes, is significantly correlated (E) with the expression of its paralog, *RPP26*, but not (F) with the expression of RPP25L itself. (H) Demonstration of functional codependency. *ITGB1* is an LSEG in the upper aerodigestive that is more essential in cells with high expression of its paralog, *ITGB4*. All *P*-values presented in the figures are adjusted for multiple tests by the Benjamini and Hochberg false discovery rate (FDR) procedure. The yellow dots (D-H plots) represent cell lines that belong to the vulnerable lineages, and blue dots are for cell lines in the non-vulnerable lineages. In D-H, locally estimated scatterplot smoothing (LOESS) curves and 95% confidence intervals are shown alongside Spearman’s ρ and the FDR value.

We tested the correlation between the dependency scores and the expression of the paralogs. For LSEGs with more than one paralog, we choose the paralog with the most significant correlation. We found that out of the 404 genes, 153 LSEGs had a significant positive correlation in the predicted direction of reduced functional redundancy (Spearman’s ρ > 0, FDR < 0.05) (see example in Figure 3D-G). This result suggests that for these 153 LSEGs, the decreased expression of the paralog in the vulnerable tissues is a likely mechanistic explanation. We then tested how many paralogs show the opposite effect of a significant positive correlation between the dependency of the paralog and the expression of the LSEGs (symmetric compensation). Out of 147 paralogs with available dependency scores, only 25 showed symmetric compensation (FDR < 0.05) (Table S2).

An additional 45 LSEGs had a significant negative correlation between the dependency scores and the expression of their paralogs (Spearman’s ρ < 0, FDR < 0.05) (see example in Figure 3H). It implies that a higher expression of the paralogs, instead of compensating for the mutations, makes the cells more vulnerable to the mutations. Contrary to the 153 LSEGs implicated in functional redundancy, the 45 LSEGs tend to be expressed preferentially in the vulnerable lineages (OR = 3.207, *P* = 0.00099). This observation is consistent with functional codependency between the two paralogs that function together in a specific lineage. We also checked whether paralogs show the opposite effect of a significant negative correlation between the dependency of the paralog and the expression of the LSEG (symmetric dependency). We found that out of the 44 paralogs with available dependency scores, only 8 showed such symmetric dependency (FDR < 0.05) (Table S2).

We explored whether LSEGs with multiple paralogs have additional significant correlations with the expression of other paralogs and if it is in the same direction as the most significant one. We observed that in most cases (141/153), when the most significant correlation with one paralog was positive (functional redundancy), all other significant correlations with the other paralogs were also positive. Similarly, when the most significant correlation was negative (functional codependency), most other significant correlations were in the same direction (37/45). These results suggest that the distinction between functional redundancy and functional codependency in most cases is a property of the LSEGs. In agreement, gene ontology (GO) enrichment analysis showed that LSEGs implicated in functional codependency were enriched with transcription regulation. In contrast, LSEGs involved in functional redundancy were enriched with general essential functions (Table S3).

Collectively, these analyses suggest that the variable expression of paralogs in different lineages may explain 15.5% of LSEGs, the majority through lineage-specific functional redundancy (12%). The analyses also reveal another relationship type between paralogs that don’t compensate but depend on each other.

### Intercellular communication may explain LSEGs in blood cells

Our analysis so far may explain 54.8% of the LSEGs (n = 698; Figure 4A). We next turned to find alternative explanations for the remaining LSEGs (n = 576). To do so, we calculated the correlation between the dependency scores of the 576 LSEGs and the gene expression of all other genes. To identify general compensation mechanisms, we choose to focus on positive correlations (Spearman’s ρ > 0, FDR < 0.05). We used a stepwise rank regression (forward selection) to identify genes with independent positive correlation (not highly correlated with other genes). 28 LSEGs didn’t have any significant positive correlation. For the remaining 548 LSEGs, the stepwise regression substantially reduced the number of positive correlations from a median of 967 correlations per LSEG (Figure 5B) to a median of 18 independent correlations (Figure 4C). Overall, there were 5822 genes with a positive independent correlation to at least one of the 548 LSEGs.

**Figure 4.**
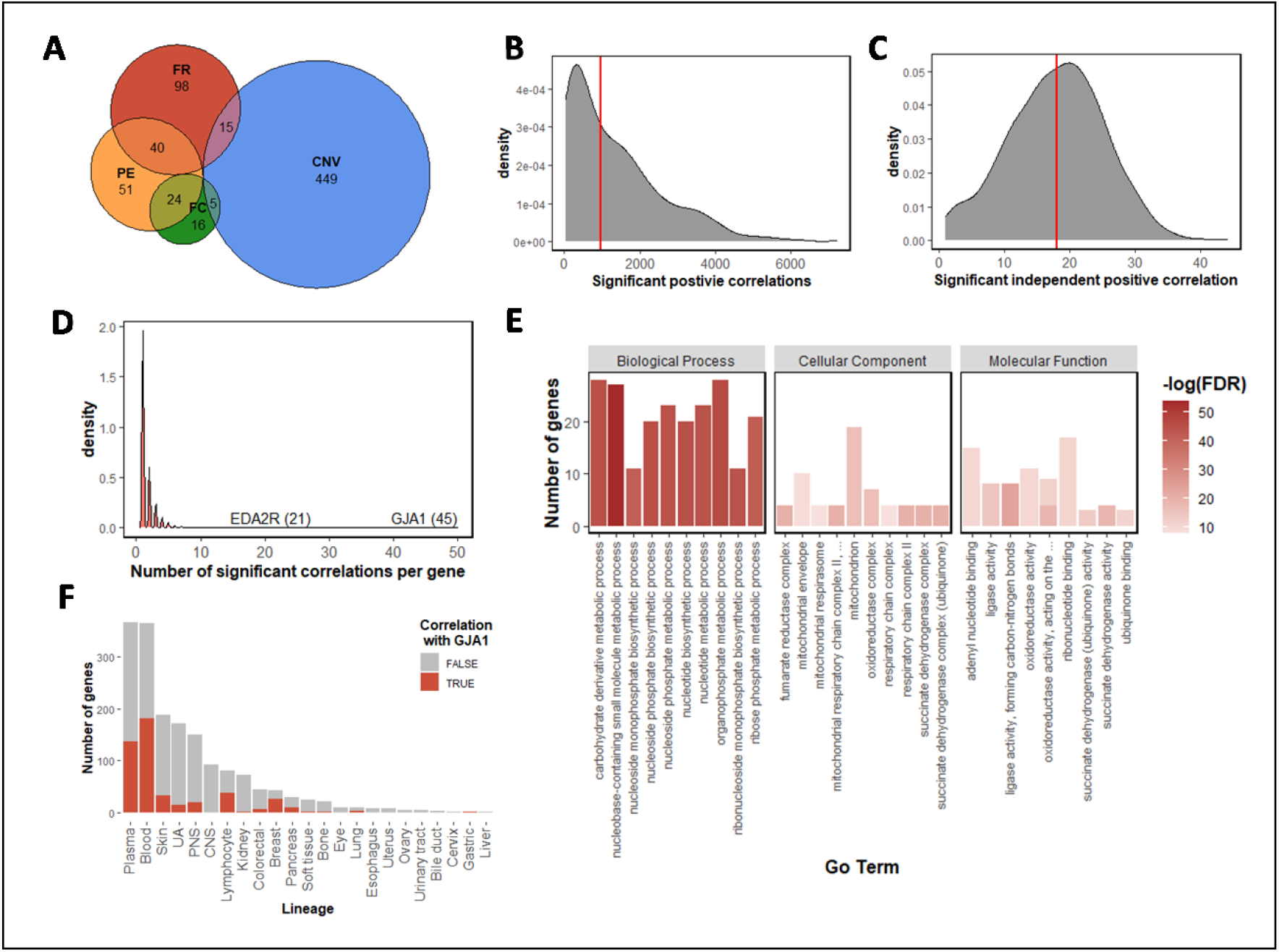
Cell to cell communication as a possible mechanism for LSEGs. (A) Euler diagram presenting the number of LSEGs that could be explained by preferential expression (‘PE’ - orange), copy number variation (‘CNV’ - blue), functional redundancy (‘FR’ - red), and functional codependency (‘FC’ - green). (B) Distribution of the number of significant positive correlations (Spearman’s ρ > 0, FDR < 0.05) for each of the 548 unexplained LSEGs. The red line represents the median number of correlations. (C) Distribution of the number of significant independent positive correlations for each of the 548 unexplained LSEGs. To identify independent correlations, we applied a stepwise forward conditional analysis. The red line represents the median number of correlations. (D) Distribution of the number of unexplained LSEGs that were significantly positively correlated with the expression of a specific gene. The names of two highly ranked genes are shown. (E) The ten most significant biological functions, cellular components, and molecular functions associated with unexplained LSEGs positively correlated with GJA1. The Y-axis represents the number of genes that overlapped a specific term. The red color intensity signifies the -log (10) of the FDR q value. (F) Comparison between the number LSEGs in each lineage that are positively correlated with GJA1 (red) or not (grey). Abbreviations: CNS – *central nervous system*, PNS – *peripheral nervous system, UA – upper aerodigestive*.

**Figure 5:**
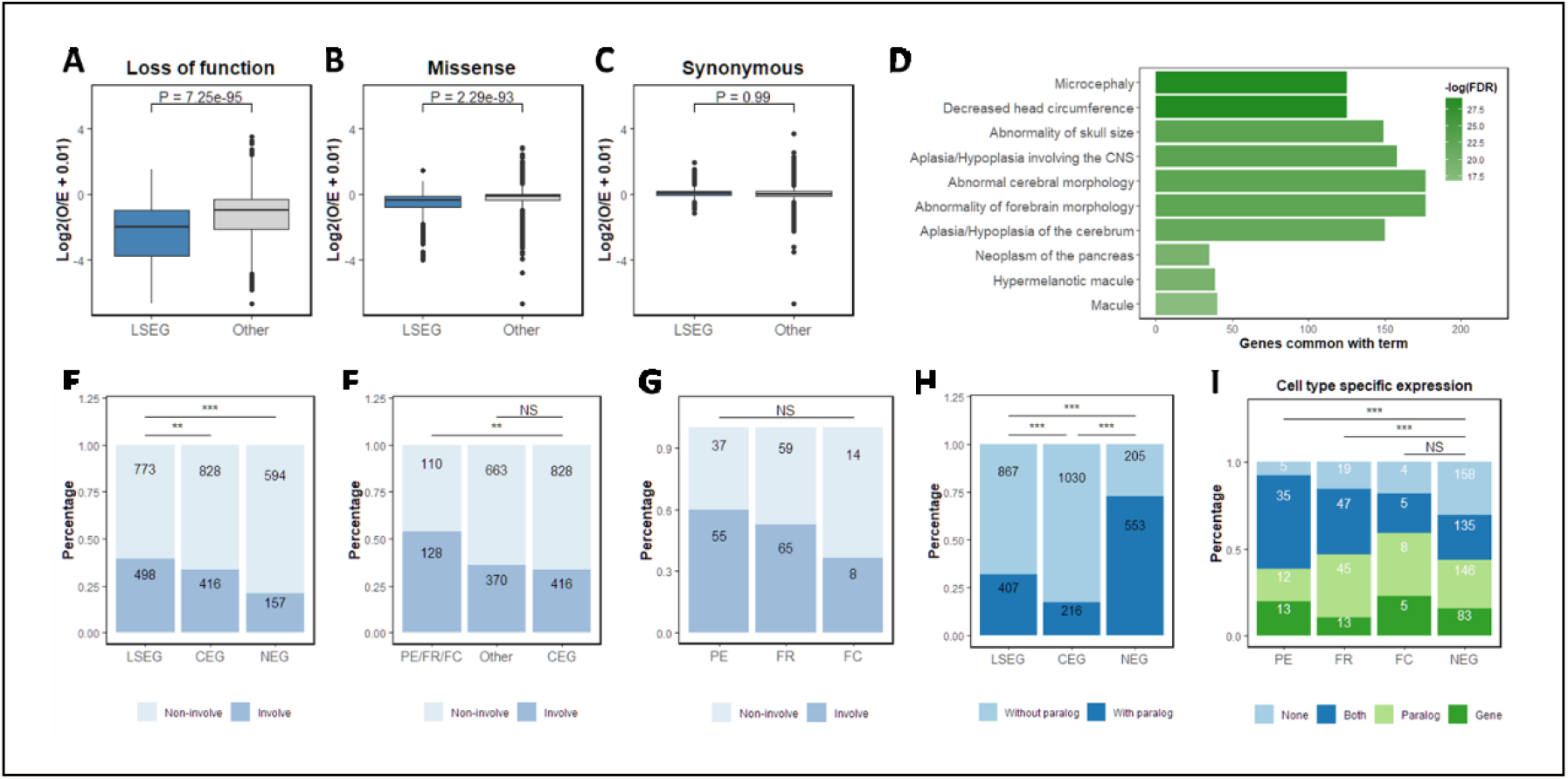
Involvement of LSEGs in human diseases and expression specificity across diverse human cell types. (A-C) Degree of intolerance to (A) loss of function mutations (B), missense mutations, and (C) synonymous mutations. Values are the Log_2_ of the ratio between the observed and expected number of mutations for LSEGs and all other genes. P-values were calculated using Wilcoxon signed-rank tests. (D) Ten most significant human phenotypes associated with LSEGs. The X-axis represents the number of genes that overlapped a specific term. Color intensity signifies the -log (10) of the FDR q value. (E-G) Percentage of genes associated with human diseases. (E) Comparing between LSEGs, common essential genes (CEGs), and nonessential genes (NEGs). (F) Comparing disease involvement between LSEGs implicated in three universal mechanisms with other LSEGs and CEGs. The three universal mechanisms, preferential expression (PE), functional redundancy (FR), and functional codependency (FC), were treated as one group. (G) Comparing disease involvement between PE, FR, and FC. (H) Proportions of genes with paralogs for LSEG, CEGs, and NEGs. (I) Number of LSEGs with cell type-specific expression for the LSEG itself (Gene), the paralog (Paralog), both the LSEG and the paralog (Both) or none of them (None). Data is shown for LSEGs implicated in PE, FR, FC, and NEGs. NS, P > 0.05; *, P < 0.05; **, P < 0.005; ***, P < 0.001.

We counted the number of LSEGs explained by each of the 5822 genes. The majority of those genes were correlated with only a few LSEGs (median = 1, mean = 1.68, sd = 1.47) (Figure 5D). We found that a particular gene, *GJA1* (also known as connexin 43), explained more LSEGs than other genes. *GJA1* was positively correlated with 45 LSEGs and was the most significant gene across the genome for 32 out of 45. *GJA1* is a component of gap junctions that allow the exchange of low molecular weight molecules between adjacent cells ^14^. Hence, these results could be interpreted as follows: mutant cells that express proteins that allow uptake through gap-junctions from surrounding cells will be less dependent on the ability to generate small essential molecules. Consistent with this hypothesis, enrichment analysis for the 45 genes correlated with *GJA1* showed that these genes are involved in the metabolism of small molecules, including nucleobase-containing small molecule metabolic process (FDR = 4.7×10^-24^), organophosphate metabolic process (FDR = 1.4×10^-20^), carbohydrate derivative metabolic process (FDR = 1.3×10^-19^), and nucleotide biosynthetic process (FDR = 1.4×10^-19^) (Figure 5E). It is important to note that the enrichment in these biological processes was highly significant compared to the level of enrichment for cellular components or molecular functions (Figure 4E).

Since gap junctions are absent in circulating blood cells, we expected the LSEGs correlated with GJA1 to be dominated by genes essential specifically in blood cells. Indeed, we found that 36 out of 45 genes were specifically essential in at least one blood-related lineage (blood, lymphocytes, and plasma cells), which was significantly more than expected by chance (OR = 4.3, *P* = 3.8×10^-5^). We also suspected that such a mechanism might explain why we had more LSEGs in tissues that lack cell-cell junctions, such as blood or plasma cells. We identified 287 LSEGs with a significant positive correlation with GJA1 expression (Spearman’s ρ > 0, FDR < 0.05). The 287 LSEGs were significantly associated with essentiality specific to blood-related lineages (OR = 9.4, *P* = 1.6×10^-46^) (Figure 4F).

### LSEGs are associated with human diseases

The LSEGs and the mechanisms we identified are based on cancer cell lines, and it raises the question of how relevant they are to human diseases in general. To test it, we first studied the overlap of LSEGs with genes intolerant to mutations. We compared the observed and expected (O/E) rate of genetic variants, which measures how strongly purifying natural selection has removed such variants from the population ^15^. Compared to other genes in the genome, LSEGs were significantly more intolerant to loss-of-function mutations (*P* = 7.2×10^-95^) and missense mutations (*P* = 2.3×10^-93^) (Figure 5A-B). As a control, we also tested synonymous mutations, which were not significantly different (*P = 0*.*99*) (Figure 5C).

Given that LSEGs are intolerant to functional mutations, we next studied the association of LSEGs with human phenotypes. Gene-set enrichment analysis (GSEA) of LSEGs revealed that these genes are enriched for developmental phenotypes, mainly of the central nervous system (Figure 5D), such as “Microcephaly” (*FDR = 2*.*2*×*10*^*-13*^) and “Decreased head circumference” (*FDR = 2*.*2*×*10*^*-13*^).

To further explore the tendency of LSEGs to be associated with human diseases, we compared the percentage of genes associated with at least one known human disease between LSEGs, common essential genes (CEGs), and nonessential genes (NEGs). We found that LSEGs are significantly more associated with human diseases compared to both common essential (*OR* = 1.2, *P* = 0.003) (Figure 5E), and nonessential genes (*OR* = 2.4, *P =* 7.6×10^-18^) (Figure 5E).

The LSEGs we identified are composed of different groups with different mechanisms. CNVs and intercellular communication are mechanisms expected to be specific to cancer cell lines or similar systems. At the same time, preferential expression, functional redundancy, and functional codependency should be important across cell types and states. Consistent with these expectations, LSEGs which were best explained by these three groups (preferential expression, functional redundancy, and functional codependency), were significantly more associated with human diseases than common essential genes (*OR =* 2.3, *P* = 5.03×10^-9^) (Figure 5F), with no significant difference between the three groups (*P* = 0.13) (Figure 5G). However, other LSEGs not belonging to the three groups were similar to common essential genes in their association to human diseases (*OR =* 1.1, *P* = 0.25) (Figure 5F). These results suggest that LSEGs are more important for human diseases than common essential genes, especially those with a widespread mechanism.

### Cell type-specific expression support the generality of the genetic mechanisms of LSEGs

Although our analysis focused on mechanisms explaining lineage-specificity in cancer cell lines, the involvement of LSEGs with human diseases suggests that the three general mechanisms we identified may be conserved in other cellular contexts. Two of the mechanisms are dependent on the expression of paralogs. When comparing the number of genes with at least one paralog, we found that 73.0% of nonessential genes have paralogs, significantly more than common essential genes (17.3%) (*OR* = 12.8, *P* = 1.0×10^-139^) (Figure 5H). The percentage of LSEGs with paralogs (31.7%) was smaller than nonessential genes (*OR* = 0.17, *P* = 1.6×10^-73^) (Figure 5H) but significantly more than common essential genes (OR = 2.2, P = 1.3×10^-17^) (Figure 5H).

The interpretation of these findings is that genes with paralogs are nonessential because of functional redundancy, while genes without paralogs tend to be common essentials. But our analysis suggests that the genes that are essential in specific cell types are preferentially expressed in those cells or have paralogs with cell type-specific expression. To test the generality of these predictions, we examined how many of the LSEGs and their paralogs expressed in specific cell types. We studied genes with at least one paralog that are best explained by preferential expression (n = 65), functional redundancy (n = 124), or functional codependency (n = 22). We compared them to nonessential genes with at least one paralog (n = 522).

The gene pairs (a gene and its paralog) can be classified into four different groups: (1) None have a cell type-specific expression, (2) Both the gene and the paralog have a specific expression, (3) Only the paralog has a specific expression, and (4) Only the gene has a specific expression. The distribution of genes in these four groups was significantly different between nonessential genes and two out of the three types of LSEGs (preferential expression *P* = 1.5×10^-6^, redundancy *P* = 3.8×10^-4^, and codependency *P =* 0.5) (Figure 5I). We directly tested the differences in cell type-specific expression frequency for the genes or their paralogs (omitting the ambiguous group when both the gene and the paralog are cell type-specific). We found that for LSEGs with preferential expression, there was a significant increase in cell type-specific expression for the gene itself compared with nonessential genes (*OR* = 2.8, *FDR* = 0.03). For LSEGs with functional redundancy, there was a significant increase in the frequency of cell type-specific expression for the paralog (*OR* = 2.3, *FDR* = 0.005). All other tests were not significant (*FDR* > 0.05).

## Discussion

We identified 1274 LSEGs that are significantly more essential in specific lineages. The 1274 genes can be attributed to different mechanisms that explain the variability between lineages. In only 9% of cases, the genes showed preferential expression in the vulnerable lineages. Our study highlights other mechanisms for the lineage-specific vulnerability to mutations, which mainly involved lineage-specific compensation. We show that LSEGs are associated with human diseases more than common essential genes. The cell-type-specific expression of the LSEGs or their paralogs is consistent with their general role in human diseases that frequently affect specific tissues.

Lineage-specific compensation is implicated in three proposed mechanisms. The first mechanism of compensation involves the gene itself (intra-allelic interaction). The compensation originates from increased gene copies, more likely to occur in specific lineages. This likely lowers the probability that mutations will disrupt all copies. Gene amplification explained a considerable proportion of LSEGs (37%). The second mechanism involves compensation by paralogs (non-allelic interactions), resulting in reduced functional redundancy in specific lineages. This proposed compensation was detected in 12% of the LSEGs (37% from the LSEGs with identified paralog). It is likely to be more widespread since redundancy exists through other mechanisms. The third mechanism is compensation between cells. Our analysis suggests that some LSEGs may arise from variation in intercellular communication that allows the transfer of small molecules to mutant cells only in specific tissues. This mechanism is relevant only in cases of genetic mosaicism when there are genetically distinct cell populations within the same individual. The metabolic cooperation through intercellular communications (such as gap junctions) was previously shown to explain the lack of phenotypes in females with X-linked diseases ^16–18^. This explains why women with a heterozygote mutation in the gene coding for hypoxanthine phosphoribosyl transferase (HPRT) have no symptoms seen in men with Lesch-Nyhan syndrome. Despite having half of their cells without the enzyme, the cells are not affected because inosinic acid, the product of HPRT, is transferred from normal to mutant cells through gap junctions. But in blood cells, compensation by other cells is not possible, and the mutant cells are negatively selected ^16^.

In contrast to the compensation mechanisms, we found that 3.5% of LSEGs (11% of LSEGs with identified paralogs) are more essential in cells where the paralog has elevated expression. The suggested mechanism is functional codependency between the paralogs. These LSEGs tend to be preferentially expressed in the vulnerable lineage, unlike the LSEGs explained by functional redundancy. We found that LSEGs with multiple paralogs in most cases were implicated in the same proposed mechanism, reinforcing the claim that there is a difference in the biological causes for variability in essentiality between these two groups. Little is known about functional codependency and its extent in humans. Previous work proposed that functional codependency stems from a direct interaction between proteins that work together as heterodimers ^19^. The assumption is that cells with higher expression of the two paralogs are more dependent on the complex and, therefore, more vulnerable to the mutation. We found that the LSEGs implicated in functional codependency are mainly involved in transcription regulation. The codependency of those transcription factors may stem from being in the same complex or having common target genes. There is already evidence of codependency for some of the genes we identified. For example, *MDM2* and *MDM4* form heterodimer required for the polyubiquitination of p53 ^20^. Other examples of pairs of paralogs known to work together are *CDC42*/*RHOJ* ^21^ and *EGFR*/*ERBB3* ^22^.

We used external datasets to show the relevance of both LSEGs and the different suggested mechanisms in human diseases and cell types. We demonstrated that these genes tend to be intolerant to missense and loss of function mutations, indicating that functional mutations in these genes are negatively selected in the human population. Additionally, we found that these genes are enriched in developmental phenotypes that involve high cell proliferation. Furthermore, we have found that LSEGs are more associated with human diseases relative to common essential and nonessential genes. The analysis of cell type-specific expression suggests that the mechanisms observed in the cancer cell lines are general and relevant to other cells and could explain their high involvement in human disease.

Our findings have important implications for studies of genotype-phenotype relationships. Studies of human diseases and potential genetic therapies for cancer are frequently based on the assumption that the cell types most affected by a mutation are those with higher gene expression. Among others, this assumption is used to connect newly identified variants to diseases or as a way to determine which tissues and developmental stages are associated with a disease. For example, in autism spectrum disorders, where the underlying brain regions and circuits are not fully understood, the analysis of genes disrupted by *de novo* mutations showed preferential expression during development in several specific brain regions ^23–25^. However, our findings indicate the need for ways to integrate the expression of the causal genes and other interacting genes to determine which tissues or cell types are likely to be involved in the disease. Thus, our results could be used to predict and explain why specific cell types are affected by a loss of a particular gene. Our work also may have specific implications for understanding the origin of lineage specificity in response to oncogenic driver mutations.

An additional and more general output of our study is that genes essential in a specific type of cells, like other forms of biological variation, can offer important insights into genetic mechanisms of genetic interactions. The mechanisms that explain the variance between cell lineages can also be relevant for the differences between human individuals with the same genetic condition and can be the basis for incomplete penetrance and variable expressivity.

## Methods

### Inferring Lineage-specific essential genes (LSEGs)

Differential dependency analysis was performed using the Limma package in R ^26^. The dependency scores were obtained from the DepMap portal (https://depmap.org/portal/) (DepMap Public 20Q3, dataset doi:10.6084/m9.figshare.12931238.v1). The dataset contains scores for 18,119 genes in 789 cancer cell lines originating from 27 different lineages. Each gene from the dataset was fitted to 27 linear models corresponding to the 27 different lineages. Each model tested for the tendency of a gene to have higher or lower dependency scores in a specific type of lineage. Moderated t-statistics, moderated F-statistic, log-odds of the differential dependency, and p-values were computed using the eBayes function from this package. The p-values were adju sted for multiple tests by a false discovery rate (FDR) Benjamini-Hochberg procedure.

LSEGs were defined as genes with significantly lower levels of dependency scores in a specific lineage compared to all other lineages (t < 0, FDR < 0.05) and with an average dependency score < -0.5. Using this threshold allowed us to exclude genes that are nonessential and, at the same time, to include genes that are not necessarily highly essential in all cell lines. LSEGs in lineages with a sample size smaller than five cell lines were excluded from the analysis. Pearson correlation test was used to test the association between the number of cell lines and the number of LSEGs per lineage. The percentage of overlap between lineages was quantified by the Jaccard Index (metric of intersection over union).

Nonessential genes (NEGs) were obtained from Hart *et al*. ^28^. Common essential genes (CEGs) were obtained from two previously reported studies, Bolmen et al.^29^ and Hart et al. ^30^, which identified essential genes for human cell lines.

### Average expression estimation in vulnerable and non-vulnerable tissues

We performed a randomized permutation test to test whether the expression of LSEGs in the vulnerable lineages tended to have more extreme values both in the positive and negative directions. We calculated for all LSEGs the mean of the absolute values of t-statistics between the expression in cell lines that belongs to the vulnerable lineages and other cell lines. This value served as the test statistic and was compared to 10,000 simulations (n) in which the lineage identity of LSEGs was randomized. We calculated the empirical P-value as (r+1)/(n+1), where r is the number of times the statistic in the simulations exceeded the true statistic ^27^. Similarly, to check whether the expression of the paralogs tends to have more extreme values (both in the positive and negative directions), we calculated for all the paralogs the mean of the absolute values of the t-statistics for the difference in expression between cell lines that belongs to the vulnerable lineages and other cell lines. This value served as a test statistic and was compared to 10,000 simulations in which the lineage identity of LSEGs was randomized. The empirical P-value was calculated as above.

### Identification of human paralogs

Human paralogs were defined using the TreeFam database (http://www.treefam.org/), classifying genes from different organisms to families based on homology. Paralogs were defined as any two human genes that belong to the same family and therefore are homologous. The number of genes with at least one paralog was compared between LSEGs, CEGs, and NEGs using Fisher’s exact test.

### Correlation tests between dependency scores, gene expression, and copy number

Spearman’s rank correlation tests were used for studying the relationship between the dependency scores of the LSEGs and the expression of 19,144 genes. Gene expression available for 783 cell lines was obtained from the Broad Institute Cancer Cell Line Encyclopedia (CCLE) expression dataset. The FDR procedure was used to adjust the *P*-values for each LSEG and across the 19,144 tests. The relationship between the copy number (obtained from the DepMap portal) and the dependency scores was tested using Spearman’s rank correlation test with FDR adjustment. Fisher’s exact test was used to test for the association between the number of LSEGs significantly associated with copy number and the direction (positive or negative) of the correlation between the expression of the LSEG and the dependency scores. Functional redundancy and functional codependency with paralogs were identified based on a significant (FDR < 0.05) positive or negative Spearman’s rank correlation between the dependency scores of the LSEGs and the expression of existing paralogs. Fisher’s exact test was used to test whether LSEGs implicated in functional codependency tend to have a preferential expression compared to LSEGs implicated in functional redundancy.

### Intolerance to mutations analysis

The overlap of LSEGs with genes intolerant to different types of mutations (loss-of-function, missense, and synonymous) was calculated using data obtained from the Genome Aggregation Database (gnomAD; https://gnomad.broadinstitute.org). For this purpose, Wilcoxon signed-rank tests were performed to compare the observed and expected (O/E) rates in LSEGs and other genes.

### Disease involvement analysis

To explore the tendency of LSEGs to be associated with human disease, we the association between genes and human diseases from the Human Protein Atlas (www.proteinatlas.org). Data were available for 1271 LSEGs, 753 NEGs, and 1246 CEGs. The number of genes involved in at least one disease was compared between different groups using Fisher’s exact tests.

### Cell type-specific expression analysis

Genes were considered with cell type-specific expression if they were defined in the protein atlas dataset to be either cell type enriched (mRNA levels in one cell type at least five times the maximum levels of all other analyzed cell types) or cell type enhanced (mRNA levels in a particular cell type at least five times average levels in all cell types)^31^.To compare between LSEGs implicated in preferential expression (PE), functional redundancy (FR), and functional codependency (FC) to NEGs, we isolated only genes with at least one paralog. We used the paralog with the most significantly correlated expression to the gene dependency scores in the case of genes with more than one paralog. We excluded genes when either they or their paralogs did not have data on cell type-specificity and cases where the gene itself had no dependency data or the paralog did not have expression data. The analysis of cell type-specific expression among the gene pairs (a gene and its paralog) was done using Fisher’s exact test.

### Stepwise rank regression models

To find alternative explanations for LSEGs, we calculated the Spearman correlation between the dependency scores of the unexplained LSEGs and gene expression of all other genes. Then, for each of those LSEGs, we created a model which included only genes with a positive spearman coefficient. We choose to focus on positive correlations to identify general compensation mechanisms. LSEGs that did not have any significant positive correlation were removed from the analysis. We used a forward stepwise rank regression procedure to identify multiple independent explanatory variables. The rank regression model is a linear model between the ranks of the LSEG dependency scores and the rank of the gene’s expression. The positively correlated genes were sequentially entered into the model, starting from the gene with the most significant positive Spearman correlation ending with the gene with the smallest correlation. The genes were considered independent explanatory variables only if their addition to the model improved it significantly (p < 0.05). The procedure stopped when there were no longer genes that could improve the model significantly. Then, to identify a general mechanism that can explain a large portion of the LSEGs, we counted how many LSEGs contained each gene in their models.

### Gene set enrichment analyses (GSEA)

The four functional enrichment analyses in the study were performed using ToppGene Suite ^32^. Using this tool, we tested LSEGs enrichment in human phenotype, LSEGs implicated in functional redundancy, LSEGs implicated in functional codependency, and LSEGs correlated with the GJA1 gene in GO terms for GO biological processes, molecular functions, and cellular components. Fisher’s exact tests were used to test LSEGs correlated with GJA1 association to blood-related lineages.

## Data Availability

This study includes no data deposited in external repositories.

